# A matrix protein acts as a cue to preserve collective motility of *B. subtilis* biofilm cells

**DOI:** 10.1101/346494

**Authors:** Nitai Steinberg, Shany Doron, Rakeshkumar Jain, Gili Rosenberg, Sven van Teeffelen, Ilana Kolodkin-Gal

## Abstract

Bacteria in nature are usually found in complex multicellular communities, termed biofilms. Biofilms are generally seen as sessile structures, resulting from downregulation of motility. However, during interspecies competition and predation, biofilm cells were shown to migrate towards competitor colonies. Here, we show that a specific extracellular matrix (ECM) protein, TasA, is essential for collective migration toward potential competitors and serves as a developmental cue that increases the formation of motile offspring from sessile chains. We reveal an effective strategy to maintain migration capacities in bacterial biofilms: besides providing a three-dimensional adhesive scaffold for the cells, TasA acts as a signal within the bacterial community.

## Introduction

Bacteria in nature are most often found in the form of multicellular aggregates commonly referred to as biofilms ^1,2^. When compared to the planktonic (free-living) state, cells in biofilms are more protected from environmental insults, including sterilizing agents, antibiotics, and the immune system. Biofilms enable bacteria to attach more firmly to their hosts and better access to nutrients ^3^^−^^9^. In many cases, bacterial biofilms are deleterious to human health, and over 80 percent of microbial infections are attributed to biofilms ^4,10^. On the other hand, some bacteria form biofilms that are beneficial. One example is the model organism *Bacillus subtilis* – a soil bacterium colonizing plant roots and protecting the host from infections by fungi and other bacterial species ^11^^−^^14^.

One main property defining biofilms is the existence of a self-produced extracellular matrix (ECM), that surrounds and protects the cells, and makes them adhere to each other or to a surface ^15^. This feature makes bacterial biofilms an especially appealing system to study multicellular development. The ECM generates a physical connection between the biofilm cells like in higher multicellular organisms ^2,16,17^. The main components of the bacterial ECM are: extracellular polysaccharides or exopolysaccharides, proteins, nucleic acids ^15^, and biogenic minerals ^18,19^.

Various genetic analyses have provided strong evidence that biofilm exopolysaccharides play a fundamental structural role in different bacterial species, impact bacterial virulence, and promote capsule formation ^20^^−^^25^. The biofilm of *B. subtilis* contains several exopolysaccharide polymers, produced by the *epsA-O* operon, and composed of glucose, galactose and N-acetyl-galactosamine ^26,27^. Colonies of mutants in the *epsA-O* operon, and specifically the glycosyltransferase gene *epsH*, lack the exopolysaccharide component of the ECM and are featureless, as opposed to the wrinkled wild type colony ^28^.

The proteinaceous component of *B. subtilis* ECM is the protein TasA, encoded by the *tapA-sipW-tasA* operon ^29,30^. TasA forms amyloidal fibers that were characterized using electron microscopy, as well as using amyloid-specific dyes ^29,31-33^. The TasA amyloid fibers are attached to the cell wall and mediate cell-to-cell adhesion in conjunction with other extracellular components ^29,34^. Colonies formed by a *tasA* deletion mutant show defective morphology, with smaller colony size and wrinkles that are less pronounced than those of the wild-type ^31^.

Being energetically costly, ECM production mechanisms are under tight regulation and activated only when biofilm formation should commence, i.e. when cells encounter a surface or other cells that already had started to produce ECM. In *B. subtilis*, two positive feedback mechanisms lead to increased ECM production: (1) the tyrosine kinase EpsAB is specifically activated by *B. subtilis* exopolysaccharides ^35^; (2) the disruption of flagellar rotation that was suggested as a mechanosensory mechanism leading to activation of ECM production ^36,37^.

Motility, like ECM production, is also a costly process involving the production of multiple components and energy investment in flagellar rotation. Thus, the coordinated expression of flagella genes, flagella assembly and rotation, are all under tight regulation ^38^^−^^40^. Only a subpopulation of cells express flagella genes, which are regulated by the alternative sigma factor D (σ^D^) 39. Until recently ^41^, *B. subtilis,* as most bacterial species, was considered to be blocked in flagellar motility when grown on high-friction surfaces ^42,43^.

Importantly, the activation of ECM production in *B. subtilis* and other bacterial species happens simultaneously with the repression of motility. The two processes are connected by complicated mechanisms, which span various levels of regulation, creating a regulatory switch between the two states. As a result of this regulatory switch, the two states, motility and ECM production, are mutually exclusive at the single-cell level ^36,37,44-59^. In *B. subtilis*, this regulatory switch depends mainly on two master regulators that together control both motility and biofilms - SinR, the biofilm master regulator, and its homologous protein SlrR^53-58^. For simplicity, this regulatory switch is termed the motility-biofilm switch. During planktonic growth, SinR represses the expression of the ECM production operons *epsA-O* and *tapA-sipW-tasA*, as well as the expression of *slrR* ^60,61^. Once the biofilm state is induced, SinR is deactivated by SinI, resulting in activation of the ECM operons and *slrR* ^60,62,63^. In turn, SlrR binds to SinR creating a heterodimer, which shifts SinR to repress the *fla/che* operon that encodes key components of motility, including the aforementioned σ^D^, instead of repressing the ECM operons ^58^. Moreover, the SlrR-SinR complex directly represses the expression of the autolysin genes, and by that induces chaining that further enhance the formation of the biofilm ^55^. Thus, the same regulator, SinR, represses either the ECM production operons or motility, but not both. Thereof, the two states of motility and ECM production are mutually exclusive. The extensive knwoledge of the decision-making processes during the development of complex communities made *B. subtilis* an especially appealing model for multicllular behavior.

Interestengly, in a recent study, we found a role for motility in interspecies interaction between biofilms. When *Bacillus simplex*, was inoculated next to *B. subtilis*, the growing biofilm of *B. subtilis* engulfed the *B. simplex* colony and eventually eradicated it. This engulfment was found to depend on flagellar motility ^41^. Moreover, because the mechanism by which *B. subtilis* killed *B. simplex* involved molecules with poor diffusion, *B. simplex* survived better when inoculated next to immotile mutant that was unable to engulf it, than when inoculated next to wild-type ^41^. This motility-dependent engulfment occurred in conditions in which flagellated motility was presumably impossible, due to high-friction forces of the surface ^42,43^. Yet, motility determined the outcome of the interaction between the two species in the conditions in which they formed biofilms. Thus, motility has a function in biofilm state that may be crucial for survival and to successful competition with other bacterial species over new niches.

In this study, we wondered whether motility also had a role in the absence of competitions. We found that TasA is a novel developmental cue for sustaining motility, and determined the molecular events that preserve the motile cell subpopulation within the biofilm. Finally, we found a novel regulator of the motility switch, a two-component system that is potentially involved in TasA sensing. Our results imply that similarly to higher multicellular organisms ^64^^−^^66^, ECM proteins can serve as signals during bacterial development, and thereby sustain collective migration.

## Results

### Flagellar motility enables biofilm engulfment of obstacles on solid surfaces

Although *B. subtilis* was previously considered to be blocked in flagellar motility on high-friction surfaces ^42,43^, in a recent study we showed that *B. subtilis* was able to engulf and attack colonies of foreign species that were inoculated adjacent to it ^67^. This behavior was dependent on the motility machinery as both *hag*, encoding the flagellin protein that constitutes the flagellar filament, and *motAB*, the motor subunit that enables flagellar rotation, were essential for complete engulfment. Therefore, we studied the role of flagellar motility in biofilm development on high-friction surfaces, and compared the development of colonies of the wild-type strain and of mutants in motility (*hag* and *motAB*) or in the main two-component system that regulates bacterial chemotaxis (*cheA* and *cheY*) (Figure 1).

**Figure 1.**
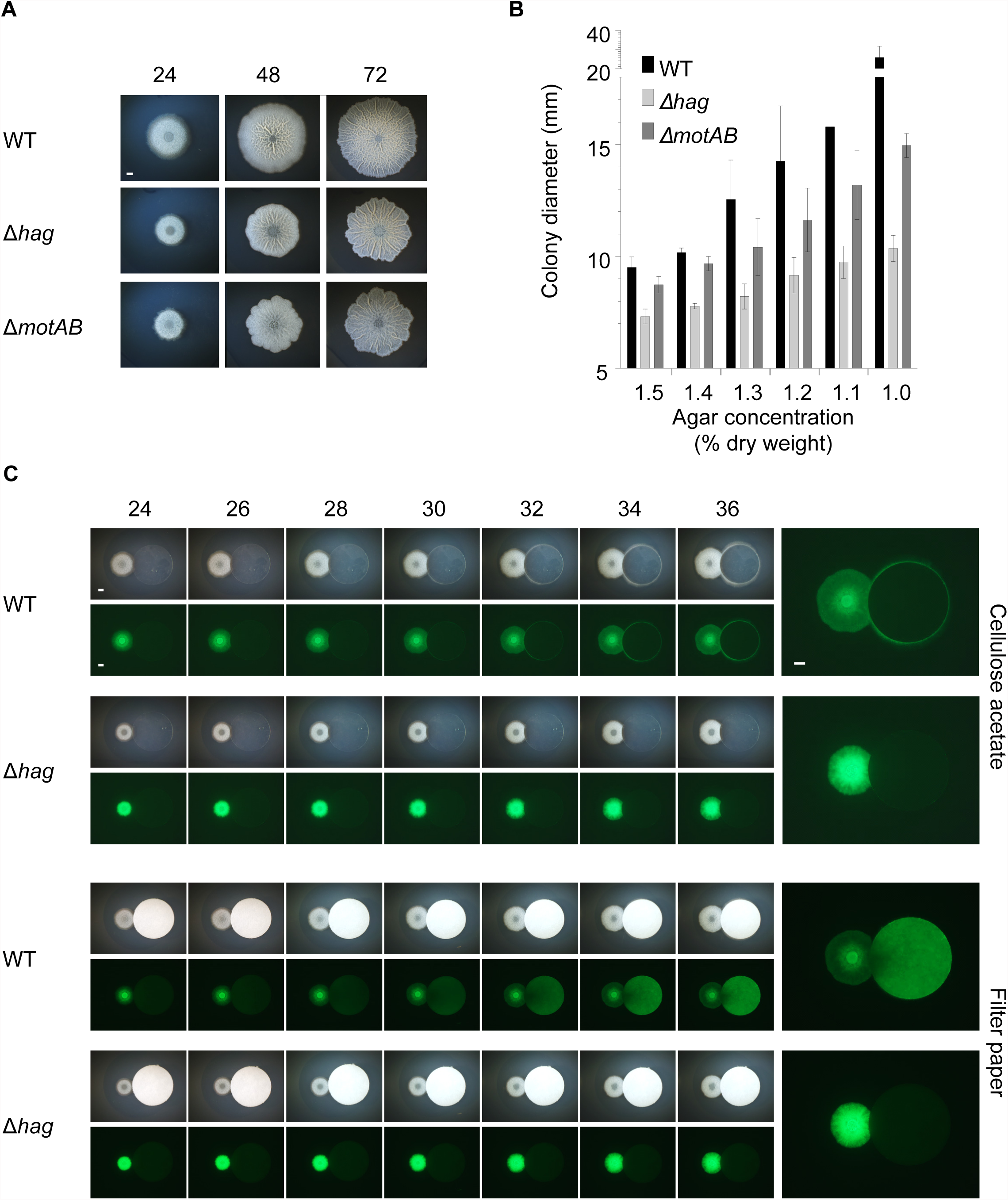
Flagellar Motility accelerates colony expansion and engulfment of foreign objects. **A** Colonies of wild-type and motility mutants grown on top of 1.5% agar plates of biofilm inducing medium (MSgg) for the indicated time. **B** Colony diameter of wild-type and motility mutant colonies grown for 24 hours on top of varying agar concentrations as indicated. Mean diameter ± standard deviation of three technical repeats is shown. Both mutants formed significantly smaller colonies than the wild-type (P-value ≤0.05) on agar concentrations of 1%, 1.3-1.5% as judged by a two tailed Student’s t-test. **C** Time lapse images of the engulfment of cellulose acetate (*top*) or filter paper (*bottom*) discs placed 0.3 cm from the inoculation point, of wild-type and Δ*haga* harboring P*_hag_-gfp* reporter, as indicated. Colonies were grown for the indicated time (in hours) on 1.5% agar MSgg plates. For each strain, bright field (*top*) and GFP fluorescence (*bottom*) images are presented. *Right –* Magnification of t=36 hours GFP fluorescence image. Scale bars represent 2 mm.

Immotile mutants formed colonies with the typical biofilm morphology (Figure 1A). However, the colony diameter was significantly smaller (Figure 1A and B). When agar concentrations were lowered, the differences between the diameters of the motility mutants and the wild-type colonies increased (Figure 1B). Thus, collective flagellar motility still exists within the biofilm enabling the expansion of the colony at a high percentage of agar, and its influence increases when agar percentage is lowered. As biofilm experiments are traditionally carried out on a high agar percentage ^68^, the residual flagellar motility may become evident only with an additional challenge. We tested whether the engulfment behavior of competing species described in Rosenberg et al can also occur in the presence of filter paper or cellulose acetate discs. The overall population was monitored by the phase (Figure 1C, upper panels), and the subpopulation expressing motility genes was examined by monitoring the cells carrying fluorescent reporter for *hag* expression (Figure 1C, bottom panels). As we found that *B. subtilis* can engulf them as well, we eliminated external influences from the foreign bacterial species. We found that cells in the engulfed area exhibit higher levels of *hag* expression, implying a higher percentage of motile cells in those areas (Figure 1C). Importantly, when a paper disc was placed near the inoculation site of wild-type *B. subtilis*, the disc became fluorescent 32 hours post-inoculation, implying that the expanded colony covered the paper disc, in contrast to the cellulose acetate disc which remained non-fluorescent.

When compared to the wild-type, motility mutants were delayed in their ability to engulf adjacent discs (Figure 1C and Supplemental Figure S1). The *hag* mutant was delayed in the engulfment of both disc types, and was unable to cover the paper disc (Figure 1C), while the *motAB* mutant was able to partially engulf both discs, but was unable to cover the paper disc (Supplemental Figure 1). It is important to note that the engulfment started more than 24 hours post-inoculation, implying that motility still significantly affected the ability of the biofilm to expand during late stages of biofilm development. Overall, we found that flagellar motility is essential for the engulfment of obstacles during late stages of biofilm development. Therefore, a substantial subpopulation of the biofilm cells must remain in the motile state in order to preserve the full capacities of the wild-type biofilm.

### The ECM serves as signal to maintain motility in the biofilm

As ECM presence was suggested to stimulate biofilm cells to turn into ECM producers ^35^^−^^37^, and as in ECM producers the motility is downregulated ^59^, we expected ECM mutants to show better engulfment capacity and higher motility gene expression. However, Δ*tasA*, a mutant in the proteinaceous component of *B. subtilis* ECM, exhibited delayed engulfment of cellulose acetate and paper discs, similarly to motility mutants (Figure 2A).

**Figure 2.**
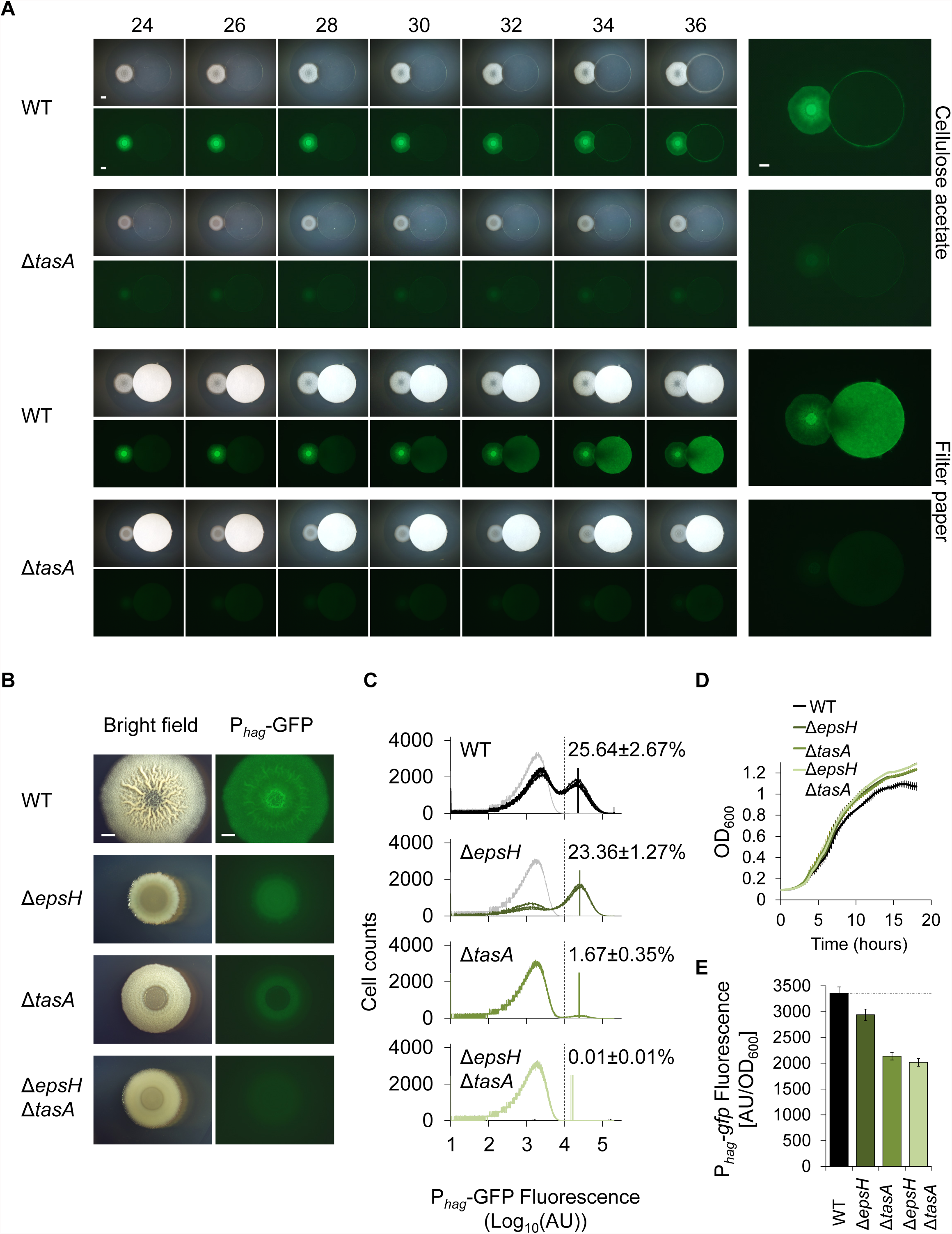
Extracellular matrix induces flagellar expression in biofilms. **A** Time lapse images of the engulfment of cellulose acetate (*top*) or filter paper (*bottom*) discs placed 0.3 cm from the inoculation point, of wild-type and Δ*tasA* harboring P*_hag_-gfp* reporter, as indicated. Colonies were grown for the indicated time (in hours) on 1.5% agar MSgg plates. For each strain, bright field (*top*) and GFP fluorescence (*bottom*) images are presented. *Right –* Magnification of t=36 hours GFP fluorescence image. **B** Colonies of wild-type and extracellular matrix (ECM) mutants harboring P*_hag_-gfp* reporter, grown on top of 1.5% agar MSgg plates for 48 hours. *Left* – bright field; *Right* – GFP fluorescence images. **C** Flow cytometry measurements of colonies such as in B. Gray – non-fluorescent control, colored – three technical repeats; Dashed vertical line – auto-fluorescence level used for gating; Solid line – median of gated population; Percentage of gated cells ± standard deviation of three technical repeats is shown. **D** Growth curves of wild-type and ECM mutants harboring P*_hag_-gfp* reporter in shaking liquid cultures. Error bars represent standard deviations. **E** P*_hag_-gfp* Fluorescence level of the cultures in D at 10 hours of growth normalized to OD level at 600 nm. Scale bars represent 2 mm. Fluorescence level of the cultures 10 hours of growth normalized to OD level at 600 nm. All matrix mutants different significantly from the wild-type (*P-value* ≤ 0.0001) as judged by a two tailed Student’s t-test.

Moreover, to our surprise, ECM mutant colonies had lower levels of flagellin expression, when compared to the wild-type colonies (Figure 2B and C). While Δ*epsH* shows only a slight reduction, Δ*tasA* shows a significantly larger reduction, with the double mutant reaching almost autofluorescence level (Figure 2B). The fact that the double mutant showed lower levels of fluorescence compared to each of the single mutants suggested separate pathways by which ECM affects motility. It is important to note that the significant reduction in flagellar expression was temporary. In later stages of biofilm development Δ*tasA* showed partial recovery in expression levels (Supplemental Figure 2).

We used flow cytometry in order to determine whether the apparent reduction is a result of reduced mean expression levels in all cells or of a reduced number of expressing cells (Figure 2C). The distribution of the wild-type clearly showed two subpopulations of cells, one that overlaps the autofluorescence levels, and one that highly expresses *hag*, in accordance with previous studies ^52^. The ECM mutants exhibited a reduction in the percentage of the *hag-*expressing cells, which supports our previous findings (Figure 2B). The median value for fluorescence intensity of the *hag* expressing cell subpopulation remained relatively similar in all strains. Thus, *hag* expression levels of the expressing cells were the same, but the number of expressing cells was lower in the ECM mutants.

To rule out that the reduction of flagellar gene expression of each of the ECM mutants was not due to a non-specific, global reduction of gene expression, we evaluated the effect of TasA on expression from different promoters that are unrelated to motility and active during biofilm formation ^69^: the promoters of the competences related gene *comG* and *OppA*. In contrast to flagellin expression, the Δ*tasA* mutant showed no reduction of P*_comGA_*-based expression compared to the wild-type (Supplemental Figures 3A and B). Similar results were observed for the expression of the transporter *oppA* (Supplemental Figure 3C). Thus, the large reduction of flagellin expression in the Δ*tasA* mutant is not due to non-specific repression.

**Figure 3.**
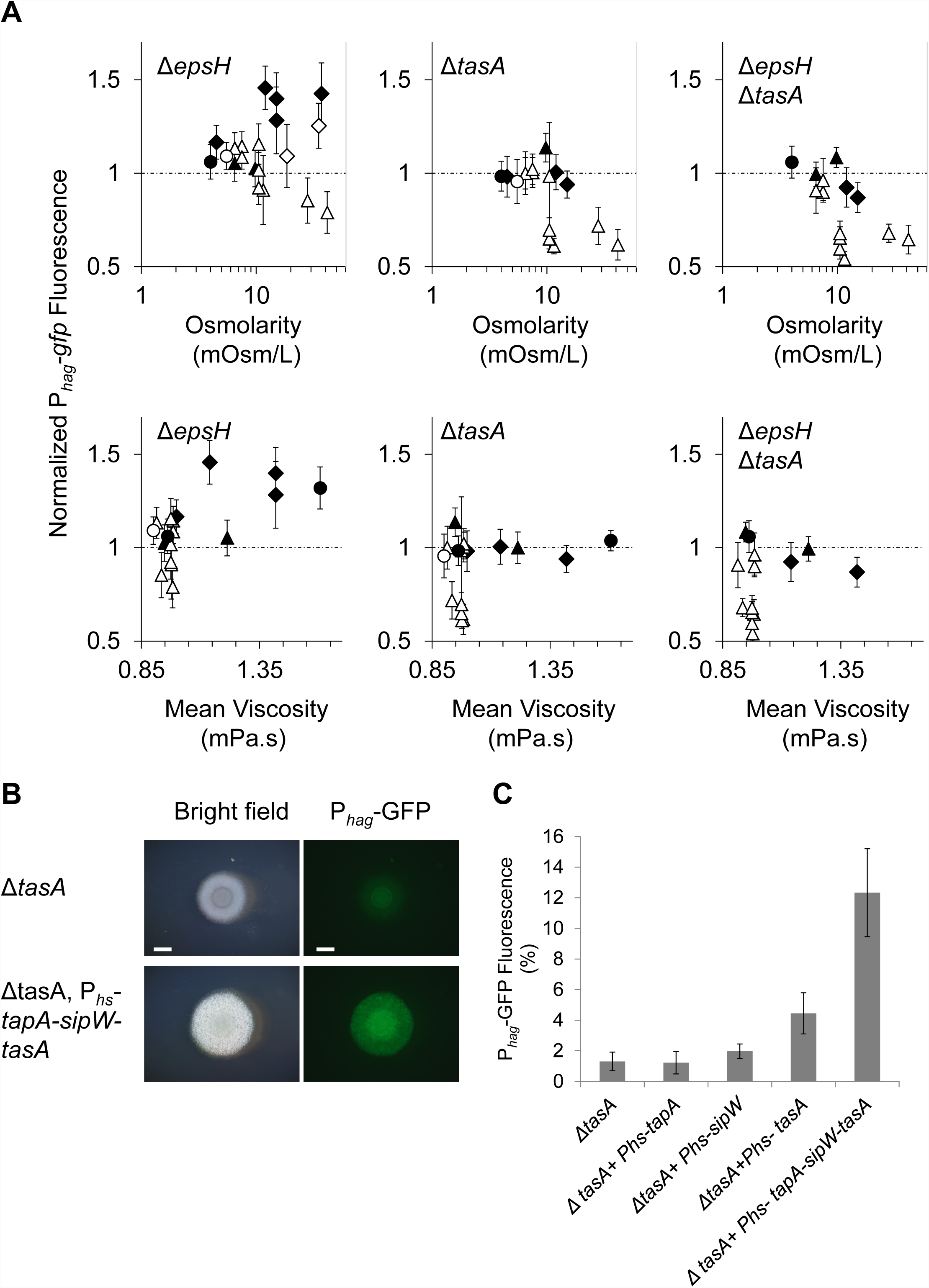
Restoration of flagellin expression levels in the ECM mutants. **A** Normalized P*_hag_-gfp* fluorescence level of the cultures treated with various substances as a function of their osmolarity (*top*) or viscosity (*bottom*). Fluorescence level at 10 hours of growth was normalized to optical density level at 600 nm, and then normalized to the level of a non-treated sample in the same experiment. Closed diamonds – Dextran 40 kDa; Open diamonds – Dextran 150 kDa; Closed triangles – Galactose; Open triangles – Amino acid mix; Closed circles – PEG 3,500 kDa; Open circles – PEG 8,000 kDa. Error bars represent standard deviation. **B** *Left –* Colonies of Δ*tasA* and Δ*tasA* harboring a *tapA-sipW-tasA* overexpression construct (P_hyperspank_-*tapA-sipW-tasA*) and a P*_hag_-gfp* reporter, grown on top of 1.5% agar MSgg plates for 24 hours. *Left* – bright field; *Right* – GFP fluorescence images. Scale bars represent 2 mm. **C** Flow cytometry measurements of colonies of the indicated strains, grown on top of 1.5% agar MSgg plates for 24 hours for 16 hours in the presence of 10 µM of IPTG. Three colonies were combined for each measurement. Percentage of gated GFP positive cells ± standard deviation of nine technical repeats is shown. Overexpression of *tasA* (P-value=0.0011), and *tapA-sipW-tasA* (P-value<0.0001) differed significantly from a *tasA* mutant as judged by a two tailed Student’s t-test.

It was shown that growth rates are different in colonies of wild-type and ECM mutants ^70^. To exclude the influence of growth rate on expression levels, we examined *hag* expression in shaking cultures, in which growth rates of the strains are comparable (Figure 2D and E). *B. subtilis* is unable to create a biofilm when grown in MSgg shaking cultures, yet it still produces ECM ^71^. Reassuringly, flagellar expression levels of shaking cultures of the wild-type and ECM mutants were consistent with those in the colonies. Thus, the reduction in flagella expressing cells was not due to reduced growth rate of the ECM mutants, nor due to different physical or chemical parameters of the developing colonies. These results further corroborate the notion that the ECM serves as a signal to retain motility.

As mentioned above, the absence of *tasA* led to a significant reduction in flagellin expression level. It was previously shown that the ECM changes physical properties of the cells’ microenvironment, such as osmolarity and viscosity, which in turn may have dramatic outcomes on biofilm development and the expression of biofilm genes ^72^^−^^74^. Therefore we asked whether the signal that is absent in a *tasA* mutant is of physical nature.

To test whether increased osmolarity or viscosity can restore flagellin expression in the *ΔtasA* strain, we added various substances with different chemical and physical properties to the different ECM mutants and examined the restoration of *hag* expression. For each added substance, we plotted *hag* expression normalized to a non-treated sample, as a function of the measured osmolarity or viscosity (Figure 3A). We chose to focus on the osmolarity range that does not cause reduction of growth rate (Supplemental Figures 4A-C). Δ*epsH* cultures showed elevation of the normalized hag expression levels when treated with artificial carbon-based polymers. There was no apparent correlation between the osmolarity or the viscosity levels and the extent of expression. In Δ*tasA* and in the double mutant, however, *hag* expression level remained the same as the non-treated sample, regardless of the added substance.

Consistent with these results, purified exopolysaccharides from wild-type pellicles restored *hag* promoter expression of Δ*epsH* to wild-type levels (Supplemental Figure 4D). Addition of purified exopolysaccharides did not affect expression level in Δ*tasA* or in the double mutant. Overall, these results suggest that the ECM exopolysaccharides, which are absent in the Δ*epsH* mutant, act as a weak chemical signal that could be complemented by a variety of substances. In comparison, Δ*tasA* seems to lack a distinct signal, indicating a link between the proteinaceous component of the ECM and flagellar motility gene expression. Reassuringly, an overexpression construct for the *tapA-sipW-tasA* operon partially restored *hag* expression levels in Δ*tasA* (Figure 3B). The restoration was dependent on the presence of the ribosome binding site within the overexpression construct (Supplementary Figure 5), implying that transcriptional regulation of motility by TasA requires the translation of the TasA protein. The expression of *hag* was restored more efficiently when TasA was expressed together with the signal peptidase SipW, and the adaptor protein TapA. In contrast, neither SipW nor TapA restored the expression of *hag* when expressed alone (Figure 3C). These results suggest that TasA has to be secreted and functional to activate motility.

As mentioned above, it was suggested that a positive feedback loop leads ECM-exposed cells to become ECM producers, turning the population as a whole to ECM production during the creation of the biofilm ^35^^−^^37^. In contrast, our findings indicate that ECM protein gene could also take part in the preservation of the important motile subpopulation in the biofilm.

### TasA stimulates reversal to motility after entering the biofilm state through the motility-biofilm switch

We found that a secreted ECM protein *tasA* has a part in the maintenance of motility in the biofilm, and that its effect is specific (Figures 2 and 3). Surprisingly, purified TasA could not restore motility to the *tasA* mutant (Supplementary Figure 6), indicating that TasA could act on a very local scale or to serve as a cell autonomous developmental cue. Importantly co-culturing of the *tasA* mutant with a *tasA* overexpressing strain partially resorted the expression of flagellin by four folds. To determine which molecular processes are affected by TasA, and how they increase the motile cell subpopulation, we decided to move to single-cell analysis using fluorescence microscopy. To that end, we compared wild-type and Δ*tasA* cells, harboring P*_hag-_gfp* as a reporter for motility and P*_tapA_-cfp* as a reporter for ECM production. These strains were grown in MSgg shaking cultures and placed on agar pads for time-lapse microscopy (Figure 4, and Supplemental Videos S1 and S2).

**Figure 4.**
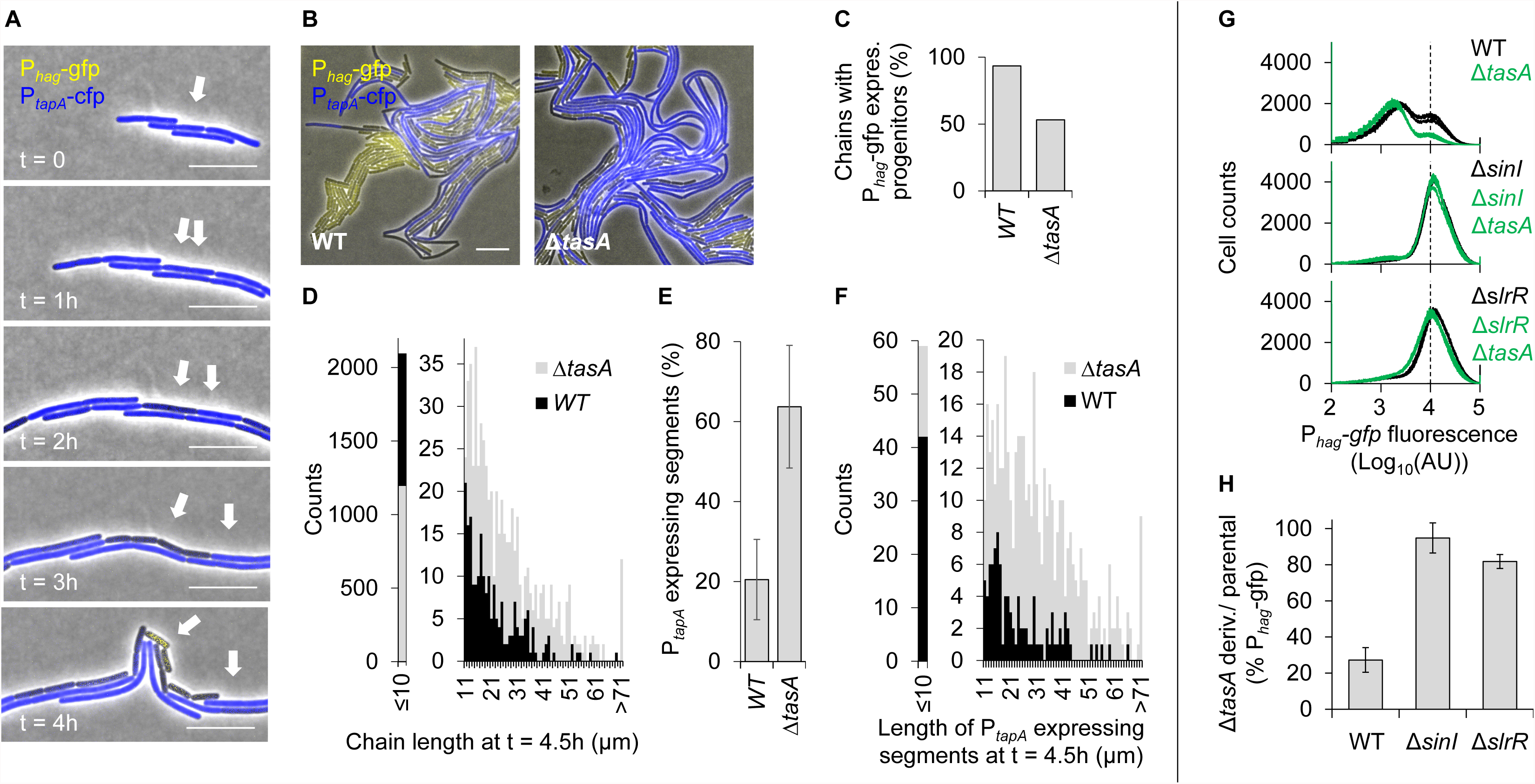
TasA stimulates reverting to motility after entering the biofilm state. **A** and **B** Cells of P*_tap_A-cfp*, P*_hag_-gfp* expressing WT and Δ*tasA*. Images were taken on agar MSgg pads (See Methods); GFP fluorescence channel – yellow, CFP fluorescence channel – blue. A Images were taken at the indicated time after inoculation. B Images were taken 7 hours after inoculation. **C** Fraction of chains for which at least one of their progenitors activated motility during the time course (4.5 hours). Number of chains with motile progenies in *ΔtasA* was significantly smaller than the WT (Student’s t-test, *P-value* ≤ 0.0001). **D** Distribution of cell and chain length at 4.5 hours. **E** Fraction of total length of all cells expressing P*_tapA_-cfp* at 4.5 hours. Chains of Δ*tasA* were significantly longer 1 hour before activation of motility compared to WT (Student’s t-test, *P-value* ≤ 0.0001). **F** Length distribution of segments in individual cells, chains or part of chains, in which P*_tapA_* is expressed. Chain length is the length of the cell centerline between the two cell poles. Chains were counted manually. **G** Shown are flow cytometry measurements of colonies of the indicated genetic background harboring P*_hag_-gfp* reporter, grown on top of 1.5% agar MSgg plates for 48 hours before harvest. Black – *top* – wild-type, *middle* – Δ*sinI*, *bottom* – Δ*slrR*. Green – *top* – Δ*tasA*, *middle* – Δ*sinI* Δ*tasA*, *bottom* – Δ*slrR* Δ*tasA*. Three technical repeats are shown for each strain; Dashed vertical line – auto-fluorescence level used for gating; **H** Ratio of the fractions of P*_hag_-gfp* expressing cells (above autofluorescence level) between: (1) wild-type and Δ*tasA*; (2) Δ*sinI* and Δ*sinI* Δ*tasA*; (3) Δ*slrR* and Δ*slrR* Δ*tasA*. Error bars represent standard deviation calculated according to propagation of error, of three technical repeats of each strain. Scale bars represent 10 µm. See Methods for parameter definitions of the microscopy analysis. See also Supplemental Videos S1 and S2.

As expected, cells were either expressing the motility reporter, the ECM reporter, or none, but never both at the same time (Figure 4A). ECM*-*expressing cells were usually part of a chain, while motility-expressing cells were individual cells. This is consistent with previous reports that autolysins are activated together with the motility regulon by σ^D^ to break the septum between cells inside chains ^75-^^80^. Interestingly, quite frequently, cells in the chain stopped expressing the ECM operon, and the progeny of these cells activated motility expression after a few cell divisions. In order to further explore the relation between TasA protein and the expression of motility, we examined cells harboring partially functional TasA-mCherry fusion proteins, together with the P*_hag_-gfp* reporter (Supplemental Figure 7), in a strain where the native TasA allele was deleted. Similarly to the P*_tapA_-cfp* results, cells inside chains often stopped expressing TasA-mCherry. During a few cell divisions TasA-mCherry amounts decreased, and the progeny of those cells activated motility. These findings are in agreement with the known motility-biofilm switch, and motility and biofilm states being mutually exclusive. Furthermore, they indicate that during the period, in which TasA affects the biofilm-motility switch, TasA most likely acts in proximity to, or directly on its producer.

Using P*_tapA_-cfp* and P*_hag_-gfp* transcriptional reporters, we measured fluorescence intensity and the distribution of cell lengths at different time points after the transition to the agar pad (Figure 4B-F and Supplemental Video S2). To study the effect of TasA on the biofilm-motility switch we focused our analysis on cells that were chained and expressed the P*_tapA_-cfp* reporter at the beginning of the experiment, and on their progeny during the 4.5-hour long time lapse. We counted similar numbers of chains for the wild-type and Δ*tasA* strains at the beginning of the time lapse. With similar doubling times of 1.15 ± 0.07 hours and 1.16 ± 0.04 hours for wild-type and Δ*tasA*, respectively, we followed cells for about 4 generations of growth. Here, we used the total length of all cells and chains in the field of view as an approximation of cell number, as it is difficult to identify single cells inside a chain with confidence.

When we examined activation of P*_hag-_gfp* expression in growing chains, we noticed that for most chains of the wild-type (93.5%, 72/77 analyzed chains) at least one of their progeny cells activated motility during the experiment (Figure 4B and C, and Supplemental Video S2). In sharp contrast, for the Δ*tasA* mutant this was the case in only about half of the analyzed chains (53.2%, 42/79 analyzed chains). Moreover, the final length distribution of the Δ*tasA* mutant showed a lower number of single cells (identified as chains of length ≤ 10 µm), and overall longer chains (Figure 4D, Median: 4.3 µm for the wild-type, 6.1 µm for Δ*tasA*), reflecting a lower number of chain-breaking events. Consistently, a much larger fraction of the cells expressed the P*_tapA_-cfp* reporter (Figure 4E). This is in agreement with flow-cytometry measurements we performed using strains harboring a P*_tapA_-gfp* reporter showing a higher percentage of P*_tapA_* expressing cells in the Δ*tasA* mutant (Supplemental Figure 8A and B). The longer chains in Δ*tasA* were the ones that expressed P*_tapA_-cfp* (Figure 5F), excluding a direct effect of TasA (expressed from the *tapA* upstream promoter) on autolysin activity.

**Figure 5.**
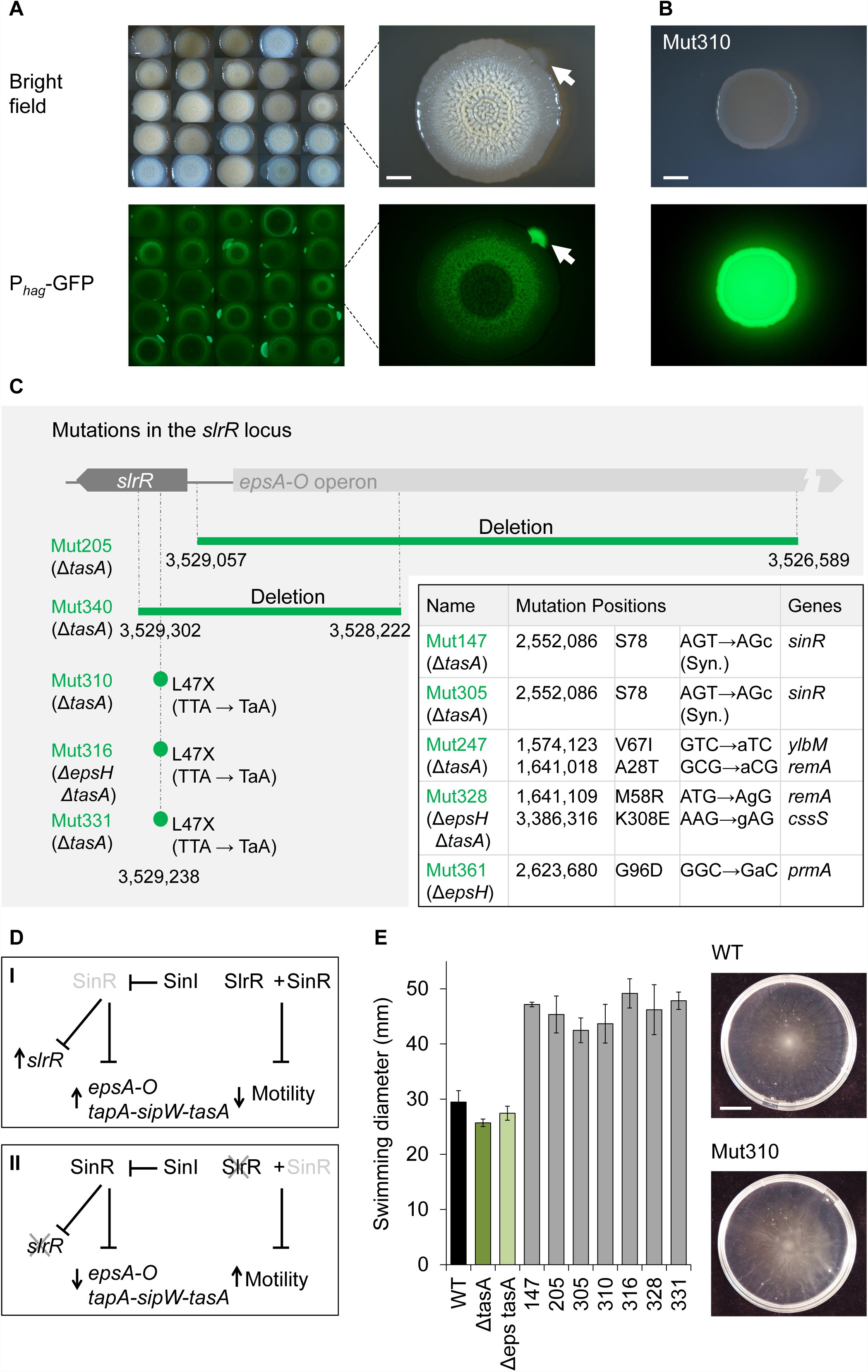
Spontaneous hypermotile mutants appear in Δ*tasA* colonies and target motility-biofilm switch regulators. **A** *Left* Overview of 25 representative Δ*tasA* colonies harboring P*_hag_-gfp* reporter and grown for 5 days and more, showing fluorescent protrusions at the colony’s edge. *Right* – Enlargement of a Δ*tasA* colony and a typical protrusion marked with a white arrow. *Top* – Bright field, *Bottom* – GFP fluorescence images. Scale bars represent 2 mm. **B** An example of a colony formed by mutants isolated from protrusions such as in A. *Top* – bright field, *Bottom* – GFP fluorescence images. Scale bars represent 2 mm. **C** *Gray frame* – Mutations in the *slrR* locus that were found in isolated mutant strains as in A. *Table* – additional mutations found in isolated mutant strains as in A: Green – strain name, in parenthesis – ancestor strain. **D** Schematic model of SinR/SlrR regulation on biofilm and motility genes; (I) During planktonic growth the expression of both ECM operons and of *slrR* is repressed by SinR; During the transition to the biofilm state, SinI represses SinR, leading to the activation of the ECM operons and *slrR*. In turn, SlrR binds to SinR and leads it to repress the motility genes; (II) In *slrR* mutant, as in the isolates found in this screen, SinR no longer represses the motility genes, thus motility is unregulated, and SinR returns to repress the ECM operons. **E** Diameter of swimming ring in MSgg 0.25% agar after 10 hours of growth at 30°C of the indicated strains, shown are mean ± standard deviation of three technical repeats. **F** Image of the swimming plates of wild-type and one of the mutants in A. Scale bar represents 2 cm.

Overall, our time-resolved analysis of the reversion from the biofilm state to motility revealed that TasA promotes a switch from the biofilm state back to motility. In the absence of *tasA,* the switching rate from biofilm to motility is decreased and cells tend to stay longer in the biofilm state.

We thus hypothesized that TasA acts through the motility-biofilm switch. To further test this hypothesis, we examined the influence of deletion of known regulators of the motility-biofilm switch on the effect of TasA on motility. We tested whether upon deletion of these regulators TasA will still have an effect on reverting from the biofilm state to motility. As expected, deletion of either *sinI* or *slrR,* the master regulators controlling the motility-biofilm switch, resulted in higher numbers of motile cells (Figure 4G), but reassuringly, the *tasA* deletion did not further affect the number of motile cells (Figure 4G and H). In order to further test the possibility that *tasA* acts specifically upstream of the SinI/SlrR switch, we examined a deletion mutant in *degU*, a global regulator that induces matrix production independently of SinI/SlrR ^48^. In the Δ*degU* genetic background, *tasA* deletion still caused a reduction in the number of motile cells (Supporting Figure S9A and B).

In conclusion, we showed that the ECM protein TasA acts as a developmental cue that increases the switching rate of cells that are in the biofilm state back to motility, and it does so through the motility-biofilm regulatory switch.

### The CssRS two-component system is a novel part of the motility-biofilm switch

As we showed above, motility is important in the biofilm environment, and TasA participates in the maintenance of the motile cell subpopulation via the motility-biofilm switch. In order to further corroborate that TasA acts upstream to the motility-biofilm switch, we followed Δ*tasA* colonies, harboring *P_hag_-gfp*, for the emergence of suppressor mutants in which motility level was restored. Targets of such suppressor mutations might reveal the molecular mechanism of TasA action.

When Δ*tasA* colonies were grown for 5 days and more, protrusions began to appear at the edge of the colony (Figure 5A). Many of these protrusions were highly fluorescent, indicating high expression from the *hag* promoter. Many additional mutants were isolated from the circumference of the colony, in areas that showed high florescence but did not protrude. In total, 80 hypermotile mutants were isolated from 141 Δ*tasA* colonies. Even after many passages (see Methods), the strains kept the hyperfluorescent phenotype, indicating a genetic change (Figure 5B). Using soft-agar (0.25%) swimming assay, we confirmed that at least in the representative mutants that were examined, this high expression indeed resulted in faster swimming (Figure 5E). Therefore, we termed these suppressor mutants hypermotile mutants. We sequenced the genomes of 9 isolated mutants (Figure 5C). Four out of the 9 mutants contained mutations in the *slrR* locus (Figure 5C and D). One had a deletion of ∼2400 bp that included the beginning of the *slrR* locus and the *epsA-O* operon (Mut205 in Figure 5C), and three had the same mutation (L47X) leading to a premature stop codon in the *slrR* gene. Using PCR and Sanger sequencing, we identified 8 additional mutants in the *slrR* gene. Out of them, six had the L47X mutation, and one had a deletion that overlaps the same region (Mut340 in Figure 5C). Two of the isolates had a synonymous mutation in the biofilm master regulator *sinR* at a specific serine codon (Figure 5D). Synonymous mutations in serine codons of *sinR*, including the mutation that was discovered here, were found to affect levels of SinR, leading to changes in expression of the ECM genes ^81^. Our results show that changes in SinR levels affect flagellin expression, probably through its effect on the *fla/che* operon together with SlrR. In two other isolates, two different non-synonymous mutations were found in the biofilm regulator *remA*. Each of these isolates harbors a second mutation that might have a separate effect. Thus, most of the suppressor mutations appearing in the Δ*tasA* colonies were in known regulators of the motility-biofilm switch. This is in accordance with our previous findings that TasA acts through the motility-biofilm switch, thus its absence plausibly led to selection of suppressor mutations that specifically targeted the switch.

One of the mutations identified in the suppressor mutant screen was in the two-component system CssRS. Isolate Mut328 carried a non-synonymous mutation (K308E) in the histidine kinase domain of the membrane-bound histidine kinase, CssS (Supplemental Figure S10A). The CssRS two-component system was previously shown to participate in the response of *B. subtilis* to secretion stress ^82,83^, but was not thought to be connected to either matrix production or motility. As mentioned above, this suppressor mutation appeared together with many other mutations that targeted key regulators of the motility-biofilm switch. We were intrigued by the possibility that the CssRS may be a novel player in the switch. To examine this possibility, we created a deletion mutant of the *cssRS* operon and inspected the effect on expression of flagella and ECM production operons (Figure 6 and Supplemental Figure 10). Strikingly, the *cssRS* deletion led to increased *hag* expression in the mutant colonies (Figure 6A, and Supplemental Figure S10B-D). Flow cytometry analysis of the colonies showed elevation in the motile cell subpopulation. Moreover, when we examined the effect on expression of the ECM production operons, using P*_tapA_*-GFP reporter, we saw a negative effect of *cssRS* deletion on P*_tapA_* expression in colonies (Figure 6B and Supplemental Figure 10E-G). Therefore, the two component system CssRS is regulating both motility and matrix expression.

**Figure 6.**
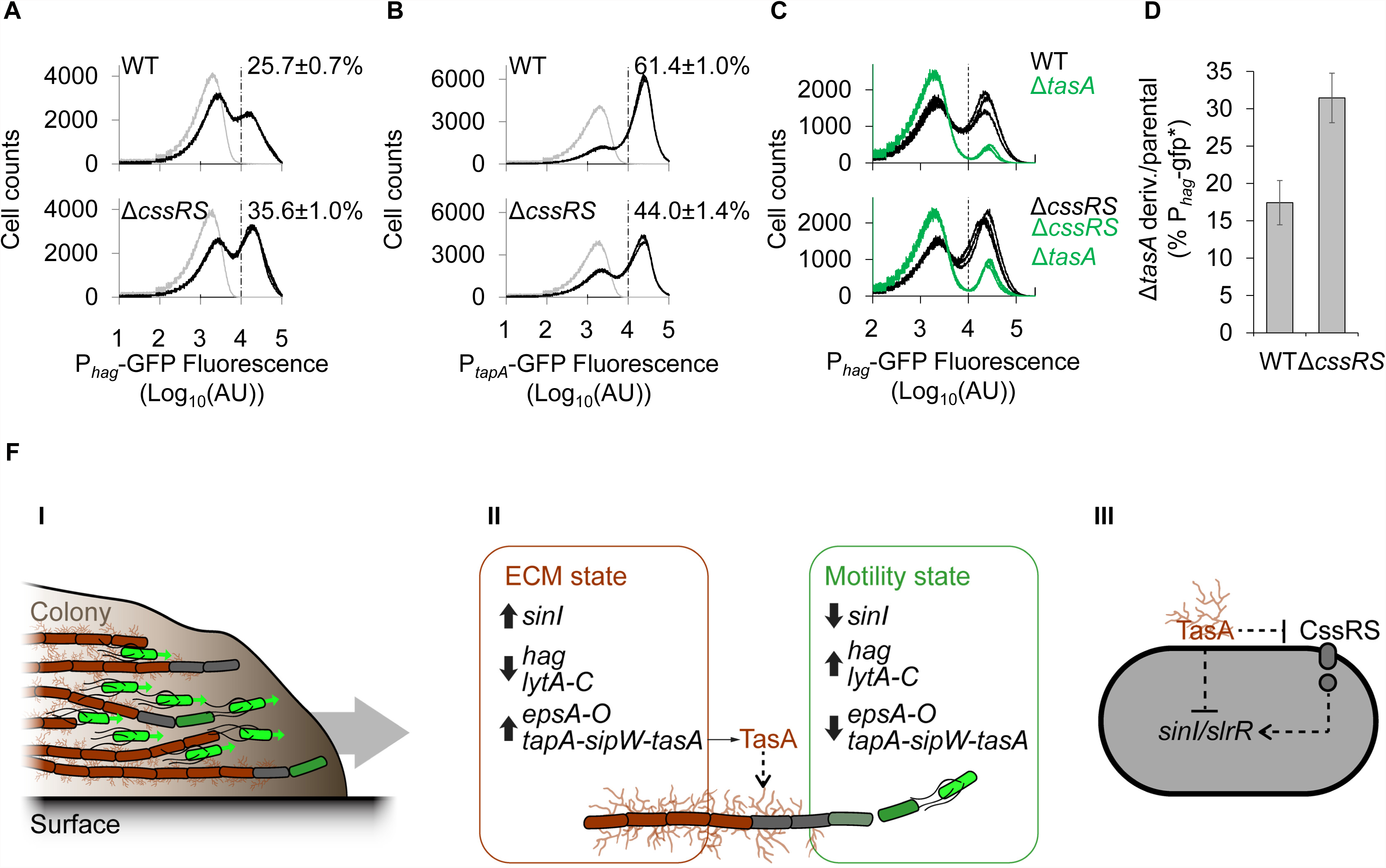
CssRS two-component system is a novel regulator of the motility-biofilm switch and is involved in TasA effect on the switch. **A** Flow cytometry measurements of colonies of wild-type and Δ*cssRS* harboring P*_hag_-gfp* reporter, grown for 24 hours on top of 1.5% agar MSgg plates. Gray – non-fluorescent control, black – three technical repeats; Dashed vertical line – autofluorescence level used for gating; Percentage of gated cells ± standard deviation of three technical repeats is shown. **B** As in A for strains harboring P*_tapA_*-*gfp* reporter. **C** Flow cytometry measurements of colonies of the indicated genetic background harboring P*_hag_-gfp* reporter, grown on top of 1.5% agar MSgg plates for 24 hours before harvest. Black – *top* – wild-type, *bottom* – Δ*cssRS*. Green – *top* – Δ*tasA*, *bottom* – Δ*cssRS* Δ*tasA*. Three technical repeats are shown of each strain; *dashed vertical line* – auto-fluorescence level used for gating; **D** Ratio of the fractions of P*_hag_-gfp* expressing cells (above autofluorescence level) between: (1) wild-type and Δ*tasA*; (2) Δ*cssRS* and Δ*cssRS* Δ*tasA*. Error bars represent standard deviation calculated according to propagation of error, of three technical repeats of each strain. **E** (I) Schematic representation of an expanding biofilm colony: Motile cells are represented in green, and ECM-producing cell-chains, as well as the TasA amyloidic fibers, are indicated in brown. The co-existence of both subpopulations in the biofilm is important for its ability to migrate. (II) *Top –* Simplified overview of the genes up or down-regulated in the ECM production (Brown rectangle) or the motility (Green rectangle) cellular states. *Bottom* – An ECM-producing cell-chain transitioning into motility, with TasA produced during the ECM production state stimulating this transition. (III) TasA acts upstream to the regulators of motility-biofilm switch that induce ECM production. CssRS activating ECM production and TasA may negatively regulate its activity.

Interestingly, it was suggested that a specific membrane kinase KinD is sensing the polysaccharide component of the ECM, and this signal leads to increased phosphorylation of Spo0A ^74,84^. A deletion of kinD had little or no effect on the response to TasA absence, indicating that TasA acts independently from KinD (Supplementary Figure 11). Similarly and in contrast to CssRS, a single mutation in the additional kinases that act upstream to the phosphorelay of Spo0A (KinA-C ^85^) failed to reduce the effect of TasA on motility (Data not shown). In contrast, when we examined flagellin expression in a Δ*cssRS* background, deletion of *tasA* decreased the number of motile cells, but to a significantly lesser degree than in wild-type background (Figure 6C and D). This raises the possibility that CssRS is linked to TasA sensing.

## Discussion

In this study, we describe a novel role for motility in bacterial biofilms, and we identify and characterize the events that maintain the motile subpopulation in the single cell level. It ^86^ has been postulated for a long time that during biofilm development flagellar motility has to be downregulated. Flagellar motility was thought to be important only during early stages of surface approaching and sensing, or in the very late stages, when biofilm dispersal commences ^11^. Nevertheless, flagellated cells were suggested to play alternate roles in biofilm development: In biofilms of *E. coli,* the presence of motile flagellated cells in mature biofilm was reported initially by Domka *et al.* ^86^, and these flagellated cells later demonstrated to play a role in shaping colony morphology ^87^. However, the role(s) of motile cells in collective migration of biofilm cells remained largely unknown.

In various bacterial species, numerous mechanisms by which motility is downregulated simultaneously with activation of ECM production were described, and include different levels of regulation. In *B. subtilis*, the presence of ECM was suggested to induce the transitioning of motile cells towards ECM production in a positive feedback loop ^35^. Flagella rotation was suggested to be impaired in the ECM-rich environment, and this disturbance was suggested as a mechanosensory pathway leading to activating ECM production 36,37. Furthermore, deletion of the gene encoding for the flagellin protein, *hag*, did not have a notable effect on biofilm morphology in *B. subtilis*. Nonetheless, in a recent study in our laboratory, we found that the ability to engulf foreign colonies enabled *B. subtilis* to deliver these molecules more efficiently and enhanced killing ^41^. Therefore, here we sought to better understand the role and the regulation of motility in biofilms.

A striking evidence for the ability of *B. subtilis* to move on top of high-friction surfaces was the engulfment of objects placed in proximity to the colony, in a similar manner to the engulfment of *B. simplex* colonies. A mutant in the motor subunit, *motAB*, that has intact flagella that cannot rotate, showed similar phenotypes to a mutant in the flagellar protein, *hag*. This further corroborates that flagellar motility is taking place. On a surface, three major factors restrict flagellar motility: surface friction, surface tension, and lack of water ^42^. In biofilms, exopolysaccharides can attract water, and surfactants can reduce surface friction and tension. Thus, the ability to move on top of high-friction surface might be much more common, and might include many bacterial species that are capable of forming a biofilm.

In the natural ecosystems of many bacterial species, and specifically, for *B. subtilis* in the soil, scenarios in which engulfment of obstacles could aid the bacterium in outcompeting other species for access to nutrients are probably frequent. As the majority of cells are contributing to ECM production during biofilm development ^52,85,88^, the central remaining question is how motility is maintained in a subpopulation of the biofilm cells.

We first examined whether ECM affects motility. To our surprise, we saw that mutants lacking different ECM components showed lower levels of motility. The ECM protein, TasA, had a dramatic and specific effect on motility, as the mutant showed a much lower number of motile cells. The effect was highly specific, as it did not show reduction in an unrelated genetic program (competence). This suggests that TasA could be the factor that maintains the motile cell subpopulation in the biofilm.

Single-cell analysis revealed the mechanism by which TasA preserves the motile cell subpopulation. We saw that TasA acts locally and is involved in reverting the ECM producers back into motile state. TasA stimulated the cells to switch back into motility, as Δ*tasA* cells stayed much longer in the biofilm state. The idea that motile cells emerge from preexisting chains is novel and has important implications – it can allow a more uniform distribution of the motile cells within the biofilm structure, rather than two segregated subpopulations of cells.

In a recent study, it was suggested that the entrance to the biofilm state is stochastic, while reverting into motility is timely ^89^. The timer that was suggested to be a passive production-dilution mechanism, in which the expression of transcription factors is stopped at a certain moment, and then they are diluted until reaching a low threshold that then allows motility to be re-activated. Here, we add an upstream layer of regulation, and this is the TasA protein itself: we show that it is essential for exiting the biofilm state. Thus, the frequent switching from ECM production to motility observed by Norman et al. could be explained by the presence of TasA on the growing chains.

Our findings that TasA acts upstream of the motility-biofilm switch are supported by additional evidence as follows: (1) When cells were prevented from entering the biofilm state, by deletion of motility-biofilm switch master regulators, *tasA* deletion had no additional influence on the number of motile cells; (2) Suppressor mutations that spontaneously appeared in Δ*tasA* colonies targeted known regulators of the switch.

Finally, an interesting non-synonymous mutation was found in the histidine kinase domain of the histidine kinase CssS, part of the CssRS two-component system, that was not known to be related to the motility-biofilm switch. As *cssRS* deletion affected the expression of both motility genes and ECM production, we suggest that the CssRS two-component system is a novel regulator of the switch. We also raise the possibility that CssRS is part of the sensing mechanism of TasA levels, but other mechanisms may exist.

To conclude, in this study we suggest a new notion of biofilms as motile, rather than sessile entities. Importantly, numerous pathogenic biofilm-forming species, such as *P. aeruginosa,* are considered to block flagellar motility while in a biofilm state. As shown here, collective motility may be fundamental to biofilm spreading. One troublesome scenario is that biofilms of virulent bacteria can spread to various tissues in the body, relying on collaboration between the ECM producers and flagellated motile cells. Thus, this new notion of collective motility, and how it coexists within the biofilm, may be found important for the development of novel anti-biofilm agents, targeting biofilm collective motility.

When considering a developmental model for biofilm formation, it is tempting to speculate that the bacterial ECM is involved in regulation of genetic programs in designated subpopulation of cells in the biofilm ^73,90^. It has been evident that in multicellular eukaryotes, cell migration depends on cell–ECM interactions ^64^. In this work, we describe a similar role for an ECM protein in as a deriving cue maintains an essential subset of motile cells within the biofilm population.

## Materials and Methods

### Bacterial strains and strain construction

All experiments were performed with *B. subtilis* NCIB 3610 ^28^. Laboratory strains of *B. subtilis* (PY79) and *E. coli* (DH5α) were used for cloning purposes. Lists of strains and primers are provided in Tables S1 and S2.

Transformation of *B. subtilis* PY79, with linearized polymerase chain reaction (PCR) products, was performed as previously described ^91^. Transformation into NCIB 3610 was performed as previously described ^92,93^. Briefly, linearized PCR products were first transformed into PY79, then the genomic DNA of the transformed strain was transformed to NCIB 3610. Linearized Plasmids were transformed directly to NCIB 3610. Mutants were verified with whole genome sequencing. Deletion mutations were generated by long-flanking homology PCR mutagenesis ^94^.

Plasmids for the generation of GFP reporter strains, or *tapA-sipW-tasA* overexpression construct were constructed as follows: (1) pYC121, which contains a functional GFP gene downstream to a cloning site and a chloramphenicol resistance gene ^71^ was used for the construction of *amyE*::P*_comGA_*-*gfp* and *amyE*::P*_tapA_*-*gfp* reporter strains. PCR fragments were amplified from NCIB 3610 chromosomal DNA, using primers with the suitable restriction sites for ligation into the plasmids. (2) *hag* promoter (according to ^39^) was amplified with EcoRI and HindIII restriction sites containing primers, and integrated into pYC121 upstream to GFP. The resulted plasmid was then cut with EcoRI and BamHI and integrated into pDR183 to create a *lacA*::P*_hag_*-*gfp*(*mls*) reporter strain. (3) pIK76 was created by inserting the EcoRI and BamHI fragment of pDR111, which contains the hyperspank promoter upstream to a cloning site, the *lacI* repressor gene ^39^ into the plasmid pDR183 which enables integration into the *lacA* locus ^95^. pIK76 was used for the construction of *tapA-sipW-tasA* overexpression strain. PCR fragment containing the *tapA-sipW-tasA* operon, ribosome binding sites, and SalI and SphI restriction sites, was integrated into pIK76.

The ligated plasmids were then transformed into *E. coli* DH5α and ampicillin resistant colonies were selected and confirmed by sequencing. The GFP reporter or overexpression plasmids were then integrated into the neutral *amyE* or *lacA* locus of the laboratory strain PY79 by transformation, as described above, and selected for chloramphenicol, spectinomycin, or MLS resistance. Extracted genomic DNA of the transformed strains were transformed to NCIB 3610 as described above.

### Media

The strains were routinely manipulated in LB (Difco) or MSgg medium (5 mM potassium phosphate, 100 mM MOPS pH 7, 2 mM MgCl2, 50 µM MnCl_2_, 50 µM FeCl_3_ for liquid or 125 µM FeCl_3_ for solid medium, 700 µM CaCl_2_, 1 µM ZnCl_2_, 2 µM thiamine, 0.5% glycerol, 0.5% glutamate, 50 µg ml^−1^ threonine, tryptophan and phenylalanine) ^28^. Medium was solidified with Bacto agar (Difco) 1.5%, or lower concentration when specified. MSgg plates were prepared on the day of the experiment and air dried under a laminar flow hood for 40 minutes prior to inoculation. Note that the iron concentration in the solid MSgg medium was 2.5-fold higher than the original recipe as this was found to improve overall morphology development as done by us previously ^88^.

Selective media for cloning purposes were prepared with LB or LB-agar using antibiotics at the following final concentrations: 100 µg ml^−1^ ampicillin (AG Scientific), 10 µg ml^−1^ kanamycin (AG Scientific), 10 µg ml^−1^ chloramphenicol (Amresco), 10 µg ml^−1^ tetracycline (Amresco), 100 µg ml^−1^ spectinomycin (Tivan Biotech) and 1 µg ml^−1^ erythromycin (Amresco) + 25 µg ml^−1^ lincomycin (Sigma Aldrich) for MLS.

### Growth and fluorescence measurements of shaking cultures

Cells from a single colony isolated on LB broth plates were grown to mid-logarithmic phase in a 3 ml LB broth culture (4 h at 37°C with shaking), and were diluted to reach equal optical density (OD). Cells were re-diluted 1:100 in 150 µl liquid MSgg medium per-well in a 96- well microplate (Thermo Scientific). Cells were grown with agitation at 30°C in a microplate reader (Synergy 2, BioTek), and the OD at 600 nm (OD600) and GFP fluorescence (485/20, 528/20 filter set) were measured every 15 minutes.

### Biofilm on solid surface assay

Cells from a single colony isolated on LB plates were grown to mid-logarithmic phase in a 3 ml LB broth culture (4 h at 37°C with shaking). Then, a 1 µL drop was spotted on solid MSgg medium. Plates were incubated at 30°C for the time period indicated in the legend for each figure. Images of bright field and GFP signal intensity were obtained with a Stereo Discovery V20” microscope with Objective Plan Apo S 0.5x FWD 134 mm or Apo S 1.0x FWD 60 mm (Zeiss) attached to an Axiocam camera. Data was analyzed using Axiovision suite software (Zeiss).

### Flow Cytometry

*B. subtilis* biofilms were inoculated as described above, and incubated for the time period indicated in the legend for each figure. Biofilms were then scraped from the plate surface and separated into single cells using mild sonication, as previously described ^96^^−^^101^. Samples were fixated in 4% paraformaldehyde (Electron Microscopy Sciences) and kept at 4°C until the measurement. Samples were measured using an LSR-II cytometer (Becton Dickinson, San Jose, CA, USA) operating a solid-state laser at 488 nm. GFP intensities were collected by 505 LP and 525/50 BP filters. For each sample, 10^6^ events were recorded and analyzed for GFP intensities. The autofluorescence level was determined in each experiment by measuring a biofilm sample from a non-fluorescent strain of the same genetic background. Then, the distribution of GFP intensities was determined using a custom Matlab code.

### Osmolarity and viscosity measurements

Different weights of dextran, glucose, galactose, xylose and polyethylene glycol, as indicated in the legend for each figure, were dissolved in the MSgg growth medium at the same concentrations that were used for the growth and fluorescence measurements (ranging from 0.2 wt% to 10 wt%). The osmolarity of these polymer solutions was measured using an ‘Advanced Instruments Freezing Point Osmometer’ (model 3300). The osmolarity was determined by subtracting the measured baseline corresponding to the MSgg solvent growth medium. The zero-shear viscosities of the polymer solutions were measured in a stress-controlled rheometer (TA-Instrument DHR-3) using cone and plate geometry.

### Exopolysaccharides purification

Pellicles were grown for 48 hours at 30 °C in 600 ml beakers on top of custom made nets. The floating biomass was then separated from the growth medium. Exopolysaccharides were extracted by us as following: Pellicles that had been formed in biofilm-inducing medium were collected, washed twice in phosphate-buffered saline (PBS) (137 mM NaCl, 2.7 mM KCl, 10 mM Na_2_HPO_4_, 1.8 mM KH_2_PO_4_), mildly sonicated, and then centrifuged to remove the cells. The supernatant was mixed with 5× ice-cold isopropanol and incubated overnight at 4°C. Samples were centrifuged at 8000 rpm for 10 min at 4°C. Pellets were resuspended in a digestion mix of 0.1 M MgCl_2_, 0.1 mg/mL DNase, and 0.1 mg/mL RNase; mildly sonicated; and incubated for 4 h at 37°C. Samples were extracted twice with phenol-chloroform. The aquatic fraction was dialyzed for 48 h with Slide-A-Lyzer dialysis cassettes by Thermo Fisher, 3500 MCWO, against dH_2_O. Samples were lyophilized. The remaining pellet was weighed and afterward dissolved in 500 *μ* L of dH_2_O.

### Fluorescent microscopy

Cells from a single colony isolated on LB plates were grown to mid-logarithmic phase in a 3 ml LB broth culture (4 h at 37°C with shaking).

For the TasA m-Cherry, P*_hag-_gfp* experiments, cultures were diluted 1:1000 in 16 ml MSgg medium in 60 mm Petri dishes, and grown for 24 hours at 23°C standing cultures. Then, cultures were mixed and 150 µl was loaded into CellASIC B04 microfluidic plates (ONIX Microfluidic Platform, Merck). These microfluidic plates contain different areas of different ceiling heights that are small enough to trap individual bacteria physically. The P*_hag_-gfp* expressing cells were trapped in the thinnest area of the plate (Number 5 - 0.7µm), while the chained and thicker TasA-mCherry expressing cells were trapped in wider areas (Number 1 - 2.3 and Number 2 - 1.3µm). We focused on those areas where the TasA-mCherry cell chains were trapped.

For the P*_hag_-gfp*, P*_tapA_-cfp* experiments, cultures were diluted 1:1000 in MSgg medium, and grown for 8 hours at 30°C with shaking. Then, cultures were diluted 1:10 in dW, and a 10 µl drop was placed on MSgg pads that were then cut out from a regular MSgg plates (see Media), incubated until all liquid was absorbed, covered with coverslips, and sealed completely with Valap (vaseline, lanolin, and paraffin 1:1:1[w/w/w]) to avoid evaporation). Images were collected using a Nikon Ti-E with a Hamamatsu Flash 4.0 V2 camera, using the following excitation and emission filters: GFP – Ex. 485/25, Em. 535/50; mCherry – Ex. 560/32, Em. 632/60; CFP – Ex. 434/17, Em. 470/28-25. Dichroic mirrors 69002bs, or 69008bs (Chroma), were used for GFP/mCherry, GFP/CFP imaging, respectively.

Time-courses of 4.5 hours were considered for analysis, as they allowed enough time for motility activation to occur, but cells were not overcrowding the field yet.

Analysis was performed manually, using ImageJ FIJI distribution ^102,103^. An event of motility activation in a chain was defined as one or more of its cell progeny reaching intensity levels in the GFP channel above a threshold defined by non-fluorescent cells. Cells that were motile in the beginning of the time-course were not considered for analysis. The number of chains in which an event occurred, or did not occur, during the time course was counted. Cell and chain length measurements were performed by manually segmenting the cells at the beginning and the end of the time course. P_tapA_ expression was defined as intensity levels in the CFP channel above a threshold defined by non-fluorescent cells. Length of segments in which P_tapA_ was expressed, either in cells, chains, or part of chains, were measured at t = 4.5 hours.

### Evolution experiment, whole genome sequencing and identification of mutations

Colonies of Δ*tasA* were inoculated as described above, and incubated for 5 days. Fluorescent protrusions from the edge of the colony were isolated on LB for single colonies. Liquid cultures of single colonies were frozen, re-streaked on LB, and the biofilm on solid surface assay was performed as described above for phenotype validation.

Genomes of nine representative strains, in addition to four ancestral strains, were sequenced. DNA was extracted using a DNeasy blood and tissue kit (Qiagen). Libraries were generated using a Nextera XT DNA sample preparation kit (Illumina). Sequencing of 100-bp single-read reads (50 cycles) was performed on an Illumina HiSeq 2500 sequencer (Illumina), using TruSeq rapid mode run reagent kits: two TruSeq Rapid SBS Kits HS (Illumina) and TruSeq Rapid SR Cluster Kit (Illumina).

Sequencing reads were aligned separately for each sample to the reference genome of *B. subtilis* NCIB 3610 (Genbank: NZ_CM000488) that was downloaded from NCBI. The reads were aligned using Novoalign 2.08.01 (Novocraft Technologies Sdn Bhd, http://www.novocraft.com) with the default parameters and [-r Random]. Detection of mutations (mismatches and insertions) was done by comparing the alignments of each sample to the alignments of the ancestor *B. subtilis* sample that was also sequenced. Genomic positions that consistently differed between both alignments (>70%) were recorded as mutations. Genomic positions with no aligned reads, but with aligned reads in the ancestor sample were recorded as deletions. Mutations in additional strains were identified using PCR and Sanger sequencing.

### Swimming assay

MSgg plates solidified with 0.25% agar were prepared a day before the experiment, and dried overnight at room temperature. Cells from a single colony isolated on LB plates were grown to mid-logarithmic phase in a 3 ml LB broth culture (4 h at 37°C with shaking), and diluted to reach equal OD. Then, a 1 µL drop was spotted on the center of the plate. Plates were incubated at 30°C for 10 hours. Images were obtained with a Nikon Coolpix P510 camera (NIKON). Data was analyzed using Axiovision suite software (Zeiss).

## Author Contributions

NS, GR and RJ performed experiments, NS, SD, and RJ analyzed the data, SVT and IKG contributed reagents, NS and IKG designed experiments, SVT designed microscopy experiments and image analysis, SVT, NS and IKG wrote the paper.

## Acknowledgments

We thank Dr. Shmuel Rubinstein and Dr. Siddarth Srinivasan, School of Engineering and Applied Sciences, Harvard University, for the osmolarity and viscosity measurements. We thank Dr. Enno Oldewurtel for the help with the microscopy experiment. We thank Dr. Zohar Bloom-Ackermann for the construction of a reported strain. The Kolodkin-Gal lab is supported by the Israeli Science Foundation grant number 119/16, France-Israel grant number 3-13021 to IKG and SVT, ISF I-CORE grant 152/1, and the Kekst Family Institute for Medical Genetics. IKG is a recipient of Rowland and Sylvia Career Development Chair.

